# Modeling Insights into Potential Mechanisms of Opioid-Induced Respiratory Depression within Medullary and Pontine Networks

**DOI:** 10.1101/2024.12.19.628766

**Authors:** Wendy L. Olsen, John A. Hayes, Dale Shuman, Kendall F. Morris, Varvara Folimonova, Thomas Tavera, Donald C. Bolser

## Abstract

The opioid epidemic is a pervasive health issue and continues to have a drastic impact on the healthcare system and United States. This is primarily because opioids cause respiratory suppression and the leading cause of death in opioid overdose is respiratory failure (*i*.*e*., opioid-induced respiratory depression, OIRD). Opioid administration can affect the frequency and magnitude of inspiratory motor drive by activating µ-opioid receptors that are located throughout the respiratory control network in the brainstem. This can significantly affect ventilation and blunt CO_2_ responsiveness, but the precise neural mechanisms that suppress breathing are not fully understood. Previous research has suggested that opioids affect medullary and pontine inspiratory neuron activity by disrupting upstream elements within this circuit. Inspiratory neurons within this network exhibit synchrony consistent with shared excitation from other neuron populations and recurrent mechanisms. One possible target for opioid suppression of inspiratory drive are excitatory synapses. Reduced excitability of these synaptic elements may result in disfacilitation and reduced synchrony among inspiratory neurons. Downstream effects of disfacilitation may result in abnormal output from phrenic motoneurons resulting in distressed breathing. We tested the plausibility of this hypothesis with a computational model of the respiratory network by targeting the synaptic excitability in fictive medullary and pontine populations. The synaptic conductances were systematically decreased while monitoring the overall respiratory motor pattern and aggregate firing rates of subsets of cell populations. Simulations suggest that highly selective, rather than generalized, actions of opioids on synapses within the inspiratory network may account for different observed breathing mechanics.

## INTRODUCTION

Opioids are considered the gold standard for the treatment of chronic and acute pain, when used appropriately. This is especially true in a perioperative environment (Hill and Canals 2022). However, the opioid epidemic remains a clear and increasingly pervasive health crisis in the United States (Ramirez et al. 2021). In 2019, overdose deaths increased by more than 50.6% since 2013 (Mattson et al. 2021). The primary cause of death in opioid overdose is respiratory depression (Hill and Canals 2022; Ramirez et al. 2021). Thus, research has been dedicated to uncovering the mechanisms associated with opioid-induced respiratory-depression (OIRD) and uncovering neural pathways to reverse OIRD (Bateman et al. 2021; Ramirez et al. 2021).

Individuals that abuse opioids ingest these drugs by one of several routes of administration including oral (approximately 90%), intravenous, and inhaled (Gasior et al. 2016; Nalamachu and Shah 2022). As such, these drugs reach the nervous system primarily from the vasculature and most readily pass the blood-brain barrier (Chaves et al. 2017) allowing them to reach all opioid-sensitive regions of the brain within the same time-frame.

Multiple regions within the brain and brainstem can contribute to OIRD (Bateman et al. 2021; Lalley 2003; Ramirez et al. 2021). Brainstem neurons work together to form an interconnected network that regulates breathing and airway protective behaviors in mammals (Lindsey et al. 2012). These circuits control the drive to breathe, cardiorespiratory functioning, and maintain ventilation (Segers et al. 2015). Rhythmic respiratory activity is facilitated and modified by interactions between the bilaterally distributed respiratory control network located within the ventrolateral medulla (Baekey et al. 2004) and certain regions within the pons (Levitt et al. 2015; Varga et al. 2020). This interconnectedness of medullary and pontine neurons is essential for eupneic breathing *in vivo* (Lindsey et al. 2012).

Commonly abused opioids like fentanyl alter breathing by activating μ-opioid receptors (µ-ORs) throughout the brainstem (Dahan et al. 2010). Pharmacological effects and adverse events are mediated by the presence of mu-opioid receptors (µ-ORs) that are found at multiple sites within the central and peripheral nervous system (Varga et al. 2020). In the ventrolateral medulla, the preBötzinger complex is active during inspiration and has been found to be sensitive to opioid agonists (Pattinson 2008). Previous research has studied these effects within intact animals and medullary slices. Results indicated that opioids have presynaptic and postsynaptic effects that alter the excitability of brainstem respiratory neurons (Gray et al. 1999, 2001). Lalley further investigated the effects of systemic administration of fentanyl by measuring intracellular membrane potentials of respiratory bulbospinal, vagal, and propriobulbar neurons in anesthetized and unanesthetized decerebrate cats (Lalley 2003). Lalley concluded that fentanyl had effects presynaptic to respiratory pre-motoneurons and motoneurons to depress neuronal activity. Additional rodent studies observed discrete, rather than continual, stepwise depression in phrenic output and inspiratory neuron discharges by opioids. These results are attributed to the effects on circuits upstream to inspiratory neurons within the preBötzinger complex (Janczewski and Feldman 2006; Mellen et al. 2003). For example, applying the opioid agonist DAMGO to the Kölliker-Fuse nucleus causes robust apneusis in a working heart–brainstem preparation of the rat (Levitt et al. 2015). However, how opioid agonists fully affect inspiratory neurons within the respiratory control network remains unclear. What has been implicated across several studies is that ventrolateral medullary and pontine circuitry, together, are affected by opioid administration and this in turn affects respiration (Burgraff et al. 2023; Dahan et al. 2010; Janczewski and Feldman 2006; Lalley 2003; Mellen et al. 2003).

Previous researchers have used intracellular recordings to measure membrane potentials of inspiratory neurons to better understand the inhibitory effects of opioids within the respiratory control network (Gray et al. 1999, 2001; Lalley 2003; Lalley and Mifflin 2017; Mellen et al. 2003). Local application of opioid agonists affects the somatodendritic µ-ORs on spatially confined presynaptic terminals while receptors in the broader region are left unaffected. This phenomenon can be difficult to interpret when the pontine and medullary circuitry, specifically the preBötzinger complex and the Kölliker-Fuse nucleus, reciprocally share sensory-motor information to generate inspiratory bursts and respiratory patterns (Varga et al. 2020). Recently, Chou and coworkers (Chou et al. 2024) disseminated findings from a computational model that was restricted to the preBötzinger complex that described plausible explanations for the observed variations in experimental responses to opioids. The group explained that their model accounts for the fixed and dynamic excitatory/inhibitory µ-OR+ neurons, cellular parameters, and network connections. They attribute discrete assigned randomness to these parameters within the model that influence individual nodes. This small level of difference is sufficient to introduce enough variance to explain the differences in experimental preparations. However, few other modeling studies specifically investigated the actions of opioids on the respiratory network. Most, including Chou et al. 2024, have been focused on limited portions of the respiratory network including the preBötzinger complex and other populations of the “core” respiratory network (Baertsch et al. 2021; Dhingra et al. 2025). Most notably, the “core” network typically does not include neuronal populations known to participate in the regulation of the breathing pattern *in vivo*, such as the pons (Lindsey et al. 1992; Rybak et al. 2008; Segers et al. 1987, 2008, 2015). Further, to our knowledge, no models addressing the actions of opioids have incorporated vagal afferent feedback which would be present in non-vagotomized animal models and humans exposed to these drugs.

### A joint neuronal-biomechanical computational model

Lalley and coworkers (Lalley 2003; Lalley and Mifflin 2017) have interpreted their results of suppressed breathing and disfacilitation of discharge patterns as an attenuation of presynaptic excitability within the pontomedullary circuitry. We have tested the plausibility of this hypothesis with a joint-neuromechanical model of the respiratory network (Lindsey et al. 2012; O’Connor et al. 2012). This model uses an integrate-and-fire neuronal network that drives deterministic equations that simulate human respiratory mechanics (O’Connor et al. 2012). The model incorporates neuron groups in the pons, raphe and nucleus of the tractus solitarius as well as the ventrolateral respiratory network (O’Connor et al. 2012); making it a unified representation of the known extent of the brainstem respiratory network (Lindsey et al. 2012). Further, it incorporates pulmonary volume-related feedback as well as a metric of laryngeal aperture that is derived from simulated motor drive from adductor and adductor muscle motoneurons (O’Connor et al. 2012). To address our hypothesis, we emulated a presynaptic action of opioids by systematically decreasing the excitability of synapses impinging on neurons in the model. The first aim of the current study was to systematically and individually decrease the strength of medullary inspiratory neuron connections within this joint neuronal-biomechanical model to examine the overall respiratory output. The second aim was to systematically, and individually, decrease the strength of pontine neuron connections within the same model. Model and trial specifications are described in the Materials and Methods section.

## MATERIALS AND METHODS

### Network construction

A joint neuronal-biomechanical model (Lindsey et al. 2012; O’Connor et al. 2012) was applied to test the hypothesis that decreasing the synaptic conductance of medullary and pontine neurons induces opioid-mediated respiratory breathing patterns. The neuronal network was made up of the functionally defined populations of neurons described in Table 1.

**Table 1.**
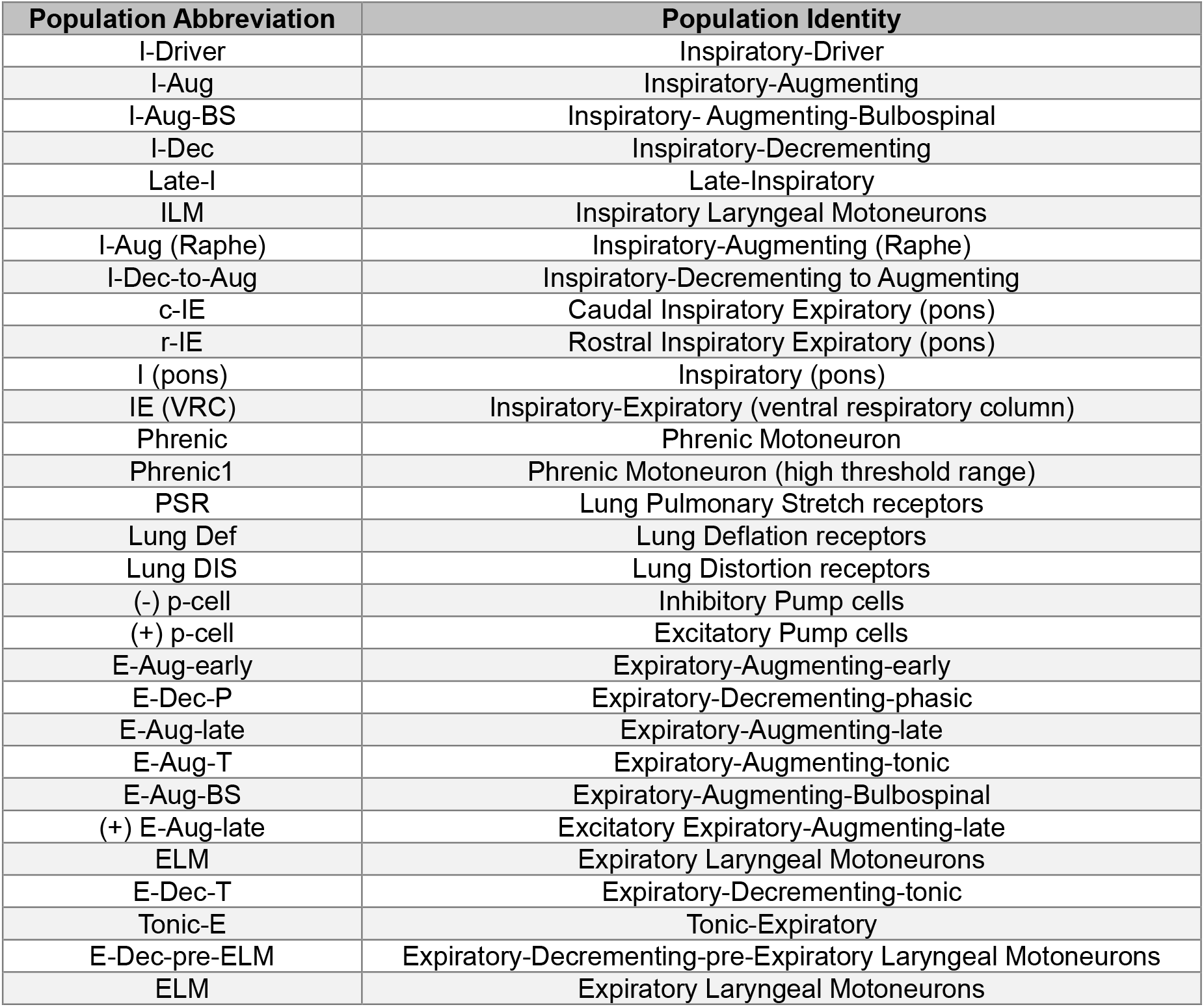
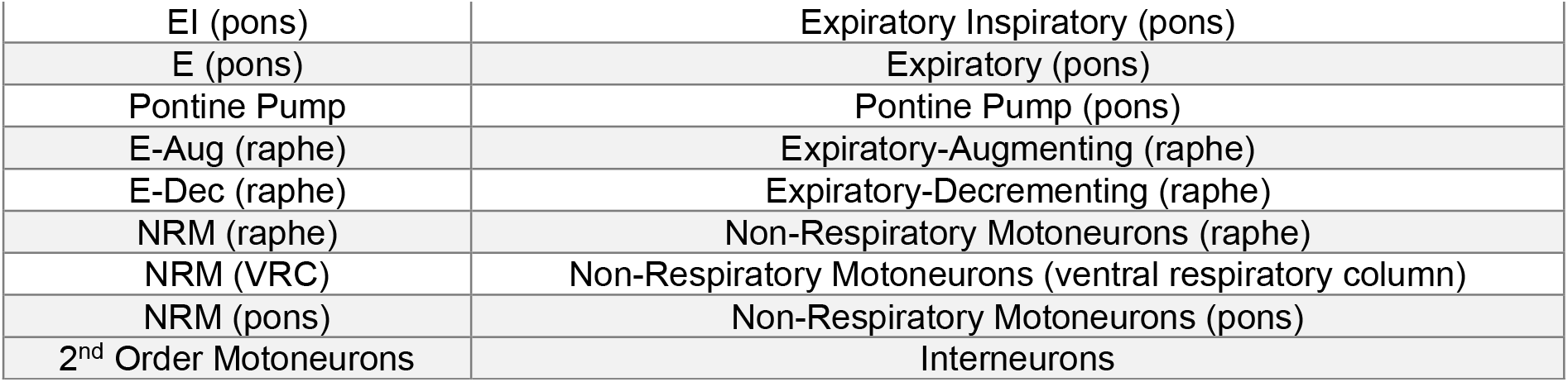
Model Neuron Populations.

#### Discrete spikes

The implementation of the neuronal components of the model come from discrete integrate-and-fire neurons where when *V*_*m*_ crosses a threshold potential (*V*_*Thresh*_) a spike state is triggered which activates downstream membrane currents of postsynaptic cells,

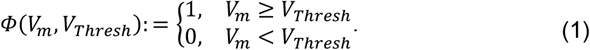

The following equations and parameters (Eq. 2-4/Table 2-3) are derived from the implementation of MacGregor’s PTNRN10 program (MacGregor 1987). Equations were integrated with a dt of 1 ms using the forward Euler method written in the C programming language.

**Table 2.** MacGregor model state variables.

| State Variable | Initial Condition | Description |
| --- | --- | --- |
| $V_m$ | -50 | membrane potential |
| $G_K$ | 0 | potassium conductance after firing |
| $V_{Thresh}$ | $V_{Thresh-0}$ | spike threshold potential |

**Table 3.** MacGregor model parameters.

| Parameter | Value | Description |
| --- | --- | --- |
| $\Delta t$ | 0.5 | time step (ms) |
| $C$ | 0.5 | accommodation parameter |
| $B$ | 5 | potassium conductance changes with C |
| $E_K$ | -10 | potassium equilibrium potential |
| $\tau_{GK}$ | 3 | potassium conductance time constant |
| $V_{Thresh-0}$ | 10 | resting spike threshold potential |
| $\tau_{mem}$ | 5 | membrane time constant |
| $\tau_{Thresh}$ | 30 | spike accommodation time constant |

#### MacGregor’s PTNRN10 model for most neurons

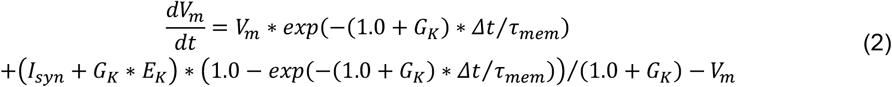

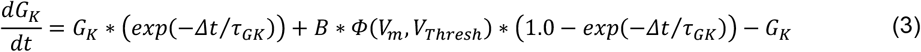

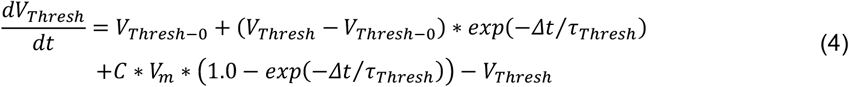

From (MacGregor 1987), when *V*_*m*_ > *V*_*Thresh*_(*h*):

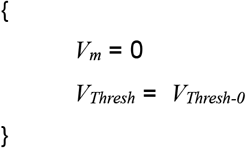

Additionally, a single population of neurons (I-Driver) were simulated using a hybridized bursting integrate-and-fire population based on Hodgkin-Huxley equations (Breen et al. 2003) described by Eq. 5-12/Table 4-5. The latter was previously developed from a continuously integrated model (Butera et al. 1999).

**Table 4.** Breen model state variables.

| State Variable | Initial Condition | Description |
| --- | --- | --- |
| $V_m$ | -50 | membrane potential |
| $h_{NaP}$ | 0.48 | NaP inactivation gating variable |

**Table 5.** Breen parameter definitions.

| Parameter | Value | Description |
| --- | --- | --- |
| $C_m$ | 21 | membrane capacitance |
| $E_{Na}$ | 50 | Nernst reversal potential for Na<sup>+</sup> |
| $E_L$ | -65 | reversal potential for leakage current |
| $\theta_{m-NaP}$ | -40 | voltage for NaP half-activation |
| $\theta_{h-NaP}$ | -48 | voltage for NaP half-inactivation |
| $\sigma_{m-NaP}$ | -6 | slope for NaP activation |
| $\sigma_{h-NaP}$ | 6 | slope for NaP inactivation |
| $\tau_{h-NaP}$ | 10000 | time constant for NaP inactivation |
| $\bar{g}_{NaP}$ | 2.8 | maximum NaP conductance |
| $\bar{g}_L$ | 2.8 | maximum leakage conductance |
| $I_{app}$ | 20 | applied stimulus current |

#### Breen model for I-Driver neurons

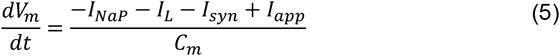

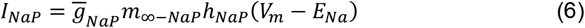

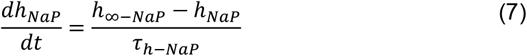

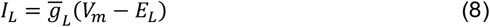

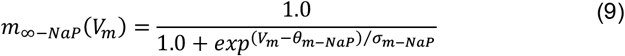

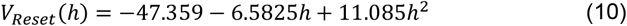

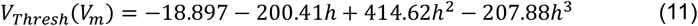

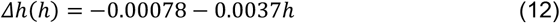

From (Breen et al. 2003), when *V*_*m*_ > *V*_*Thresh*_(*h*):

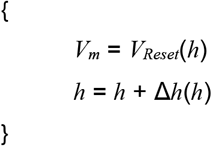

Some of the key model parameters across both models and simulation types that will be discussed are in Table 6.

**Table 6.** Key model parameters for neurons.

| Parameter | Description |
| --- | --- |
| $\bar{g}_{syn}$ | maximum synaptic conductance |
| $h_{NaP}$ | NaP inactivation |
| $I_{NaP}$ | slowly-inactivating “persistent” sodium current |
| $V_m$ | membrane voltage |
| $\tau_{h-NaP}$ | time constant of NaP inactivation |
| $I_{syn}$ | synaptic current |
| $\tau_{syn}$ | time constant for synaptic current |
| $I_{app}$ | applied stimulus current |
| $\Delta I_{app}$ | change in applied stimulus current |
| $\bar{g}_{Excit}$ | maximum excitatory conductance |
| $\bar{g}_{Inhib}$ | maximum inhibitory conductance |

#### Synaptic transmission

Communication between neurons was transmitted through the *I*_syn_ variable for both models as in equation 11,

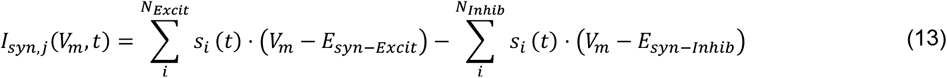

Where the *i*’s are the presynaptic connections, *j* is the postsynaptic neuron of interest, *N*_*Excit*_, the number of excitatory inputs, and *N*_*Inhib*_, the number of inhibitory inputs. Each neuron maintains a FIFO (first-in, first-out) queue matrix (*Q*_*k*_) that tracks the incoming conduction time (the columns) from each synaptic input (the rows).

The minimum and maximum conduction time is variable but across all populations the maximum is 10 ms and the minimum is 0 ms.

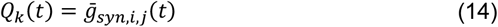

These values are then used to calculate each synaptic state variable (*s*_*i*_) representing the input coming from a specific source

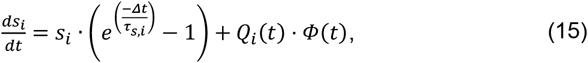

and used in the generalized *I*_*syn*_ equation 13 for the two neuronal models.

#### Biomechanical components

The present model was derived from the above equations paired with *in vivo* data that enhanced the development. Ultimately, motor output from phrenic motoneuron populations at each time step (0.5 ms) within the model are calculated by counting the total spikes from all the cells in each of two phrenic populations (*P*0 and *P*1) and dividing by the duration of a time step. Each population accounts for different inspiratory burst activity within the model. The output of the two is described by equation 16:

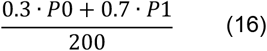

Notably, there should be a maximum value of 1 for phrenic population recruitment. This equation describes the model’s representation of inspiratory output. The lumbar motor output is handled in a similar way as the phrenic output with two lumbar motoneuronal populations (*L*0 and *L*1) as in equation 17:

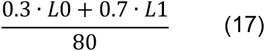

The biomechanical components of the model (Lindsey et al. 2012; O’Connor et al. 2012) were developed from transdiaphragmatic pressure and diaphragm activation while controlling the thoracoabdominal configuration (Cluzel et al. 2000; Grassino et al. 1978; Konno and Mead 1967; Song et al. 2006).

### Network simulations

The *uflsim* software package, version 1.0.36 (formally *usfsim*, Lindsey et al. 2012; O’Connor et al. 2012) utilizes a *Qt C++* cross-platform development framework written for Windows and Linux (source code may be found here: https://github.com/jahayes-ns/uflsim with Windows executable binaries that contain the “OConnor2012.snd” network file here: https://github.com/jahayes-ns/uflsim/releases/download/neuroscience/uflsim_win_1.0.36.zip).

The program includes the functionality of the program *SYSTM11* (MacGregor 1987) used in previous simulations of the respiratory network (Balis et al. 1994; Lindsey et al. 2012; O’Connor et al. 2012). The program allows neuron excitability to be modulated by injected current, and elements designated as “fiber populations” external to the network can also be used to represent transiently active afferent inputs to the network. A graphical user interface (*simbuild*) was used to modify cell parameters and network structure while the resulting model files were simulated using *simrun*. Simulations were run on 64-bit Intel-based computers under the Windows or Linux operating systems. Python scripts were developed to produce large sets of *uflsim* networks (.snd files) with varied parameters such as synaptic strengths between populations of neurons, so simulations could be executed in batches and analyzed offline with figures produced using *Matplotlib* (Hunter 2007). Network summary figures (Figs. 3D-E, 5, 6B-C, 8B-C, and 9B-C) were produced by taking the mean±SEM of relevant features from 9 distinct runs of freshly produced networks derived from a “trunk” network (Fig. 2). These 9 distinct networks were randomly generated with different random seed values.

**Figure 1.**
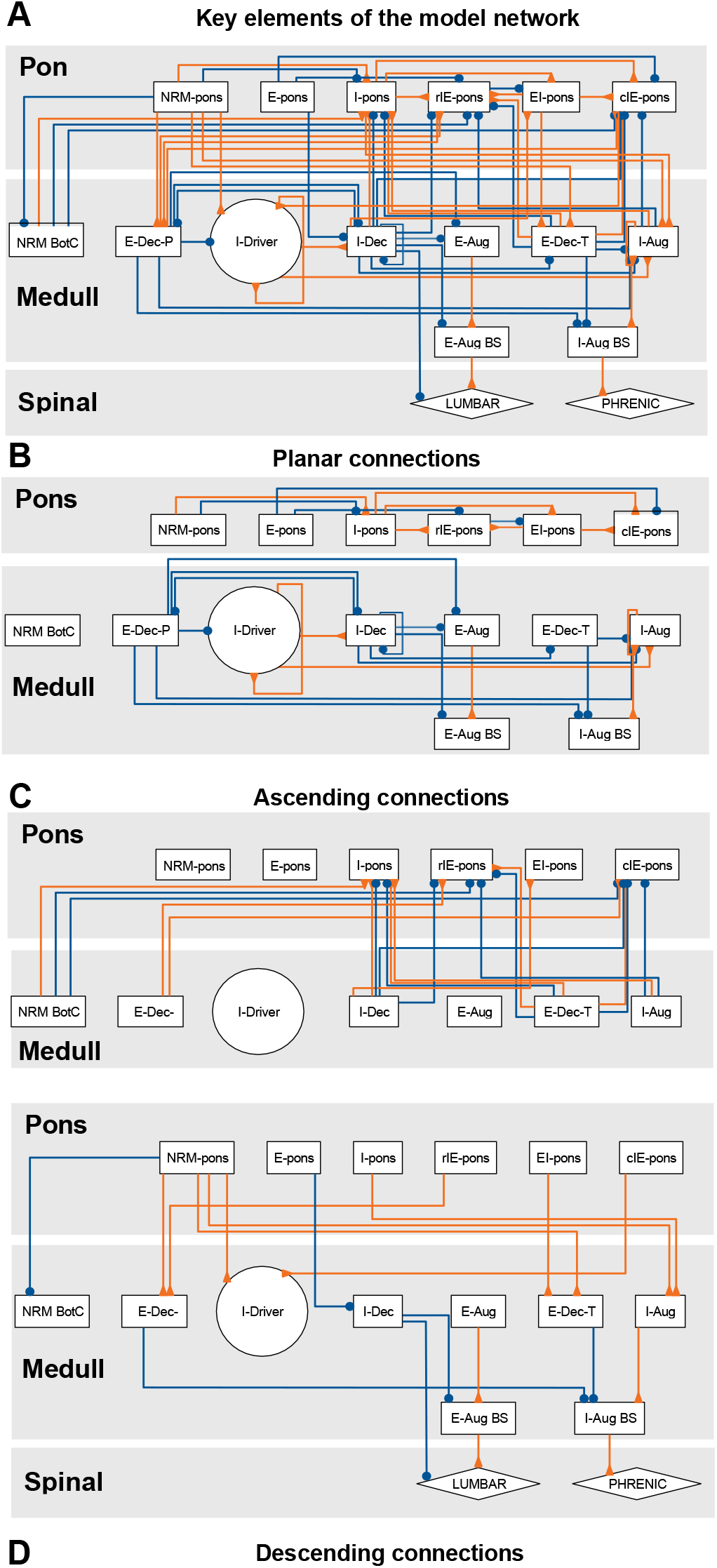
Connectivity among key elements of the model network. **A**, The core neuronal populations interrogated in this study. Orange inverted arrow connections represent excitatory connections while blue solid circle connections represent inhibitory connections. The graph is arranged roughly corresponding to hypothesized anatomical location with top-tier rostral pontine populations, middle-tier medullary populations, and spinal cord populations representing the motoneuronal output of the neuronal network at the bottom tier. **B-D**, the planar (**B**), ascending (**C**), and descending (**D**) subsets of these connections to improve clarity (O’Connor et al. 2012).

**Figure 2.**
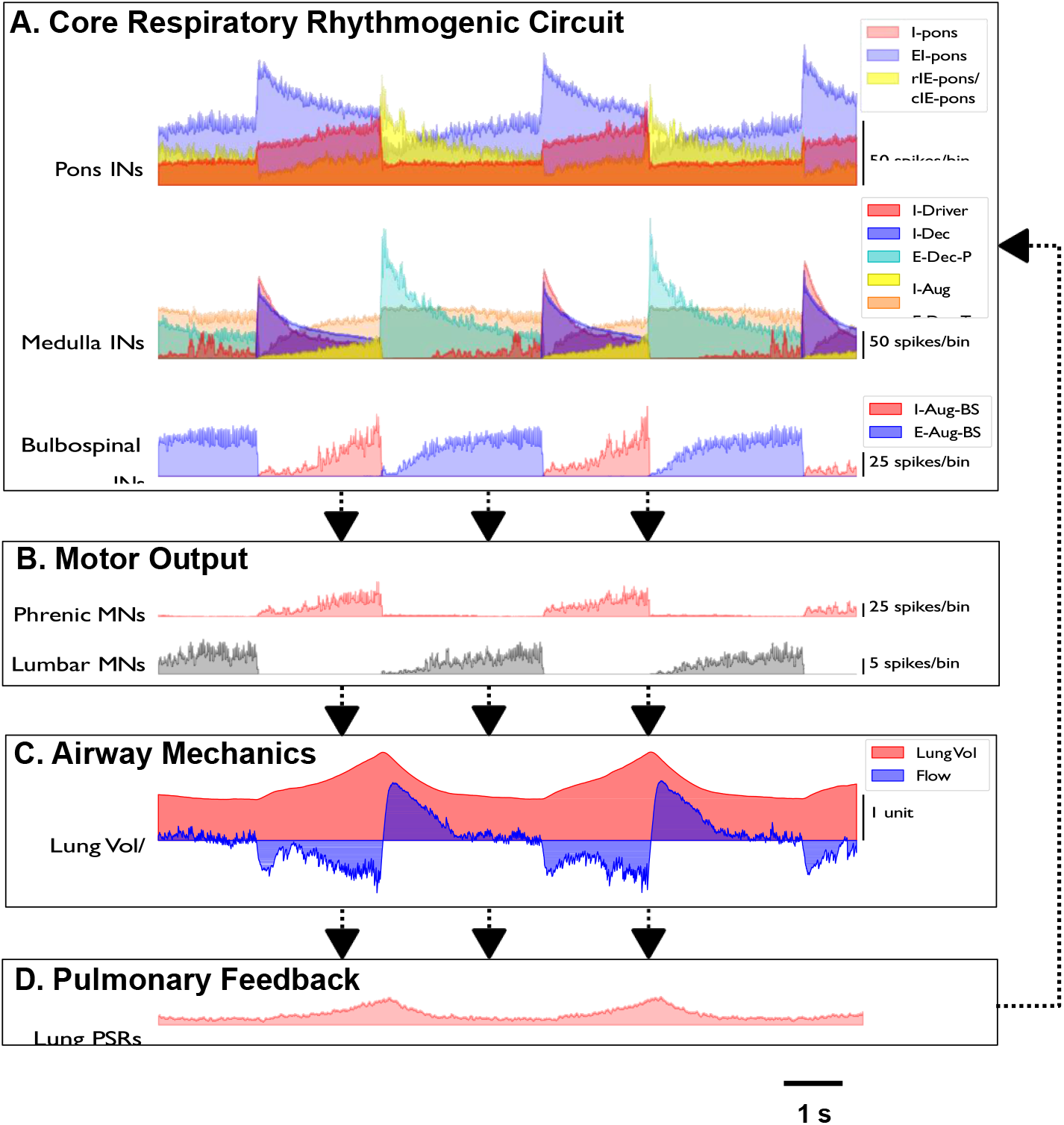
Fictive eupnea with active expiration. **A**, The core respiratory time course of activity by classes of overlayed neuronal populations. The top are pontine interneurons (Pons INs), middle medullary interneurons (Medulla INs), and bottom medullary bulbospinal premotor neurons (Bulbospinal INs). **B**, Inspiratory motor output of the simulation as expressed as phrenic motoneuronal activity (Phrenic MNs) while lumbar spinal motoneurons (Lumbar MNs) convey expiratory activity. **C**, Simulated lung volume and flow at the mouth produced by the respiratory activity. **D**, Moving average of lung pulmonary stretch receptors (Lung PSRs) activated by lung expansion. This vagal sensory information feeds back into the core respiratory circuit continuously. Arrows indicate the feedforward flow of information in the system (O’Connor et al. 2012).

**Figure 3.**
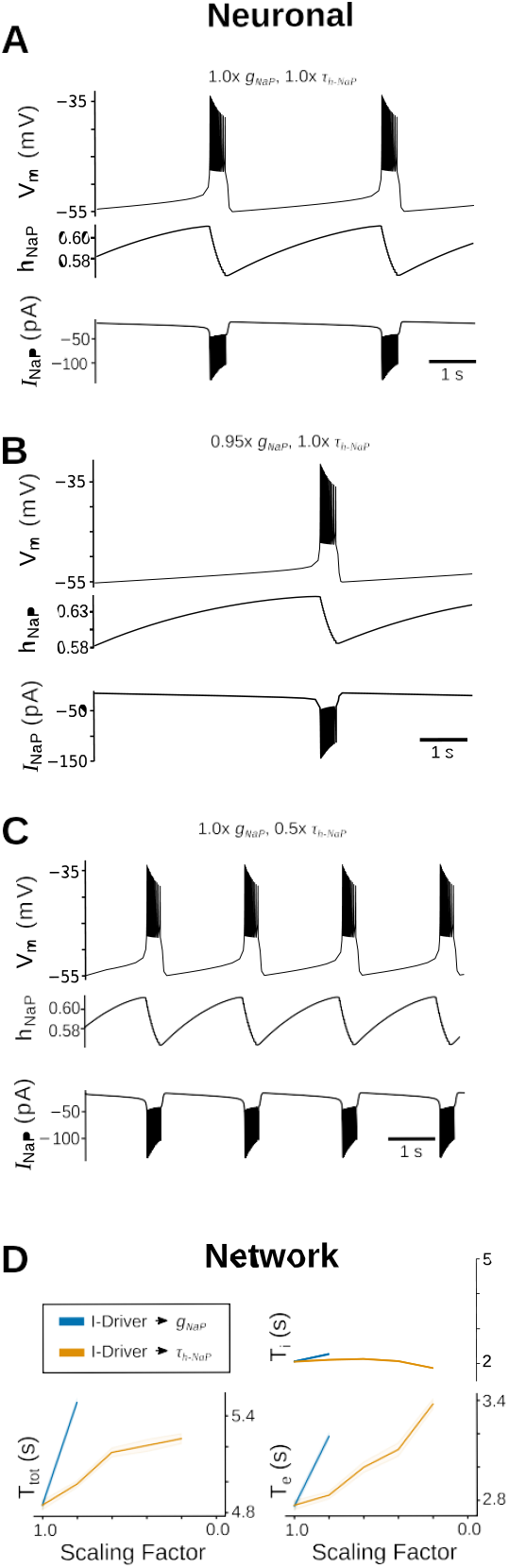
Comparison of *I*_NaP_ at the neuronal and network level. **A**, top, In a simple model of a single respiratory neuron, *I*_NaP_ can lead to periodic bursting in membrane potential (V_*m*_). (middle), The magnitude and kinetics of *I*_NaP_ are controlled by the inactivation gating variable *h*_*NaP*_ (bottom). **B**, Decreasing 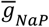 slows the burst frequency. **C**, Decreasing the rate *h*_*NaP*_ inactivates (*τ*_*h-NaP*_) increases the burst frequency. **D**, Effects on the full network period (T_tot_), burst duration (T_i_), and expiratory phase (T_e_), when decreasing 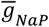 and *τ*_*h-NaP*_ in the I-Driver population of neurons. CV is the coefficient of variation. **E**, The same effects on motoneuronal output and biomechanical respiratory flow.

### Neuron simulations

Single-cell neuron simulations for Fig. 3A-C of the Breen model (Breen et al. 2003) were performed using NeuronetExperimenter (NNE, https://neuronetexp.sourceforge.net/) simulator software (Hayes et al. 2008; Song et al. 2015, 2016) with parameter values defined in Table 4-5. Panels for the figures were produced using Matplotlib (Hunter 2007).

## RESULTS

### Organization of a fictive bulbar and spinal respiratory network

We started with the network structure from a previously published respiratory model (O’Connor et al. 2012). This consists of 47 populations of neurons with up to 70-300 members each (9159 total neurons) as well as 10 fiber sources that project to these neuronal populations (1800 total fibers). Each population member had 50-200 axon terminals randomly distributed to members of its target populations (average source→target were 100.1 terminals for each pair) with 175,394,850 total terminals in the network. The identity of the neurons in each population are defined by three characteristics: 1) their typical firing phenotype under eupneic conditions, 2) their general anatomical location, and 3) their hypothesized, or experimentally identified, connectivity to other populations in the model (Fig. 1). Typically, the prefix of the eupneic inspiratory-phasing neurons is “I-” and eupneic expiratory-phasing neurons with “E-”. After this “Aug”, “Dec”, suggests the predominant discharge pattern during the respective phase as augmenting or decrementing spike rate consistent with experimental phenotypes. “NRM” indicates that a population is not eupneic respiratory modulated. Anatomical locations for the populations are sometimes specified parenthetically with “pons” or “raphe” and the remainder are by default in the medulla. The exception to the latter is the “PHRENIC” and “LUMBAR” populations of motoneurons and are meant to represent roughly the C4 and L1 levels of the spinal cord and output to muscles.

Figure 1A highlights 17 of the key populations from this model network in the context of the present study with gray boxes generally delineating the approximate anatomical location (pons, medulla, and spinal cord) for the firing phenotypes. Figure 1B shows hypothesized lateral connections within these structures (pons→pons: 6 excitatory, 4 inhibitory; medulla→medulla: 6 excitatory, 13 inhibitory), while Figure 1C and 1D show ascending (medulla→pons: 9 excitatory, 10 inhibitory) and descending (pons→medulla: 10 excitatory, 5 inhibitory; medulla→spinal cord: 2 excitatory, 1 inhibitory) connections between these structures, respectively. The complete adjacency matrix may be found in Supplemental Data S1 as both an image and a CSV file.

Figure 2 shows the population activity of each of these populations where the core respiratory rhythm-generating circuit is shown in Fig. 2A. Medullary interneurons (INs) periodically oscillate bursting between the inspiratory (I) and expiratory phases (E) with the I-Driver neurons initiating the cycles, I-Dec neurons following a similar firing pattern, and I-Aug neurons reciprocally inhibiting the others. These medullary neurons project to bulbospinal pre-motoneurons that further project to cervical motoneurons (Phrenic MNs) and lower spinal cord (Lumbar MNs) (Fig. 2B). Our biomechanical model accounts for the activity from these MNs, as well as laryngeal MNs (not shown), to model airway mechanics that drive lung inflation/deflation (Fig. 2C). Pulmonary stretch receptors (PSRs) then both excite I-Aug and inhibit I-Dec neurons during inflation closing a feedback loop (Fig. 2D).

### The relationship of I-Driver cellular properties to fictive breathing

The I-Driver neurons form the core kernel of the rhythm generator that produce the initial burst activity that percolates through the inspiratory phase (Fig. 2A). A sub-spike threshold, slowly-inactivating “persistent” sodium current (*I*_*NaP*_) produces this augmenting activity during the late-expiratory phase and is described by equation 5 (Breen et al. 2003).

Figure 3 illustrates the subthreshold activity of this *I*_NaP_ current. Fig. 3A shows the membrane voltage trajectory (*V*_*m*_) of an intrinsically bursting I-Driver-like neuron with the inactivation variable (*h*_*NaP*_) in the middle row and the *I*_*NaP*_ truncated to -50 pA in the bottom row to highlight the slowly de-inactivating current between bursts of activity. *E*_*Na*_ is the Nernst reversal potential for sodium (+50 mV) while *m*_∞_(*V*_*m*_) is the instantaneous voltage-dependent activation function for *I*_NaP_.

Figure 3B shows the same simulated neuron with the maximum synaptic conductance 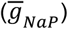 slightly decreased to a scaling factor of 95% (0.95x) which slows the bursting frequency. Further decreasing the scaling factor to 90% (0.9x) resulted in a silent neuron that relaxes to a subthreshold baseline *V*_*m*_ (not shown). In the same simulated neuron, returning to the original 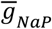 but changing the maximum time constant of NaP inactivation (τ_*h-NaP*_) to 50% (0.5x) results in a dramatic increase in the bursting frequency in Fig. 3C.

For comparison to the more expansive network model, similar graded adjustments on 300 I-Driver neurons from scaling factors 1.0x to 0.0x to 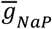 and τ_*h-NaP*_ led to changes in T_i_ (inspiratory phase duration), T_e_ (expiratory phase duration), and T_tot_ (the sum of T_i_ and T_e_, or full cycle period) and is shown in Fig. 3D. Similar to the neuron model, changing 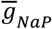 led to cessation of rhythm at relatively high levels of scaling factor for 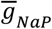 suggesting the importance of subthreshold *I*_*NaP*_ in this model to initiate the population burst activity. Remarkably, in contrast to the single neuron model, decreasing τ_*h-NaP*_ increased both the expiratory phase (inter-burst interval) and consequently the full cycle period (T_tot_) while having only a modest effect on T_i_. This qualitatively shows the dramatic impact network connectivity and synaptic properties can have on the overall production of rhythmic behavior in this model brainstem, and we explore this in more detail below.

### Generalized mechanisms for µ-OR-agonist influence on the neural control of respiration

In this study, we analyzed several distinct schemes by which µ-OR agonists may influence respiratory activity (Fig. 4). The first are comprised of cellular effects that are conceptualized as affecting baseline membrane properties through K^+^-dominated leak channels and will be examined more closely associated with Fig. 5. The key take-away from this mechanism is that, in the absence of active membrane properties more dramatic than spike-generating currents, it would simply affect the presynaptic spike rates of neurons and can be simulated by an adjustment in applied stimulus current (*I*_app_) (Fig. 4A). In contrast, mechanisms that influence connectivity strength could act through pre- or postsynaptic mechanisms (Fig. 4B) and will be considered in the subsequent Results sections (Fig. 6-10). For the purposes of this study, they are effectively the same mechanism and result in decreased *I*_syn_ (synaptic current) given a uniform spike-rate between the two.

**Figure 4.**
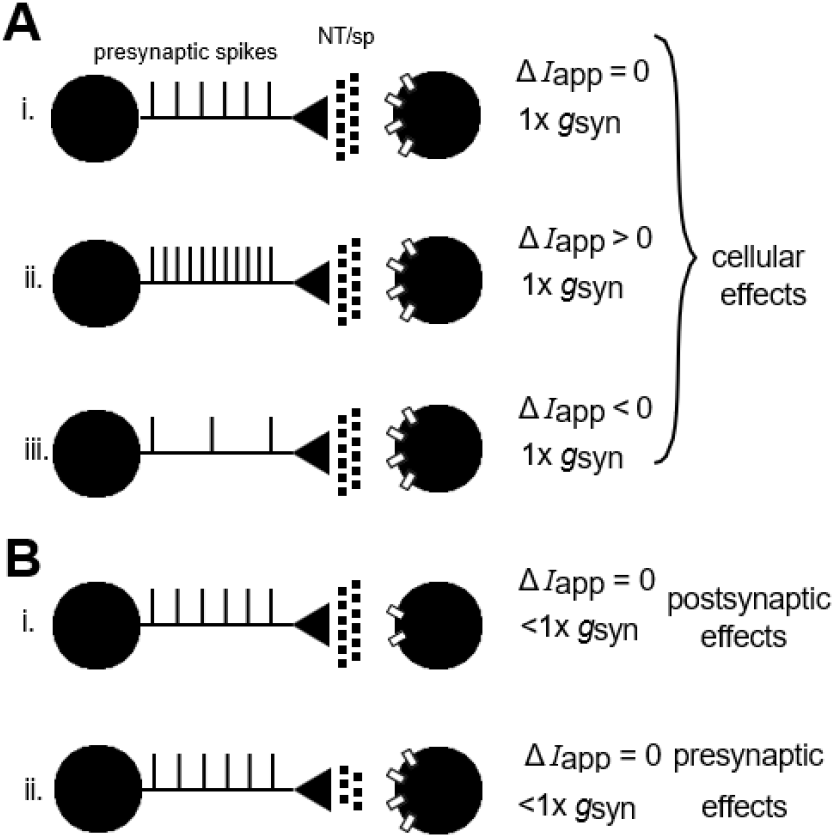
High-level comparison of possible cellular and synaptic effects of µOR-agonists. Presynaptic neuronal (left spheres) spiking with neurotransmitter release (small circles) onto postsynaptic neurons (right spheres). Clear boxes on the postsynaptic neurons represent transmitter receptors. **A**, i.), Normal presynaptic spiking activity is equivalent to normal synaptic strength (1.0x *ḡ*_*syn*_) and Δ*I*_app_ = 0. ii.), Heightened presynaptic spiking activity is equivalent to normal synaptic strength and Δ*I*_app_ > 0. iii.), Lower presynaptic spiking activity is equivalent to normal synaptic strength and Δ*I*_app_ < 0. **B**, i.), Normal presynaptic spiking activity with normal neurotransmitter release but fewer postsynaptic receptor targets is equivalent to Δ*I*_app_ = 0 and <1.0x *ḡ*_*syn*_ but weaker synaptic strength from a “postsynaptic effect”. ii.), Normal presynaptic spiking activity with decrease in neurotransmitter release but normal receptor targets is also equivalent to Δ*I*_app_ = 0 and <1.0x *ḡ*_*syn*_ but weaker synaptic strength from a “presynaptic effect”.

**Figure 5.**
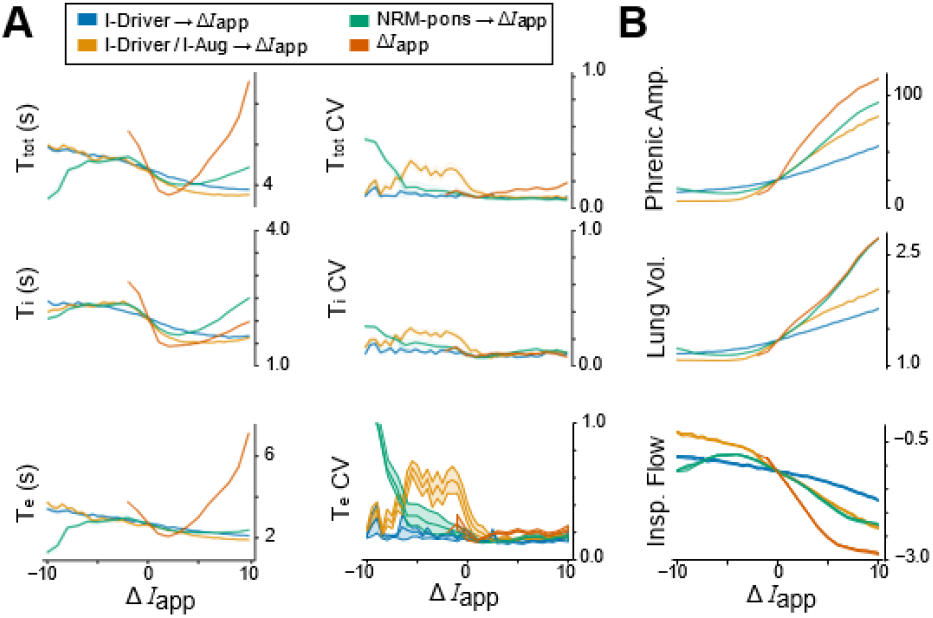
Influence of Δ*I*_app_ on network and biomechanical behaviors. **A**, T_tot_ is total respiratory period, T_i_, inspiratory phase duration, and T_e_, the expiratory phase duration. CV is the coefficient of variation for the respective quantities. **B**, (top) Maximum phrenic motoneuron amplitude during the inspiratory phase. (middle) Maximum lung volume during the inspiratory phase. (bottom) Inspiratory flow. Δ*I*_app_ is in units of pA.

**Figure 6.**
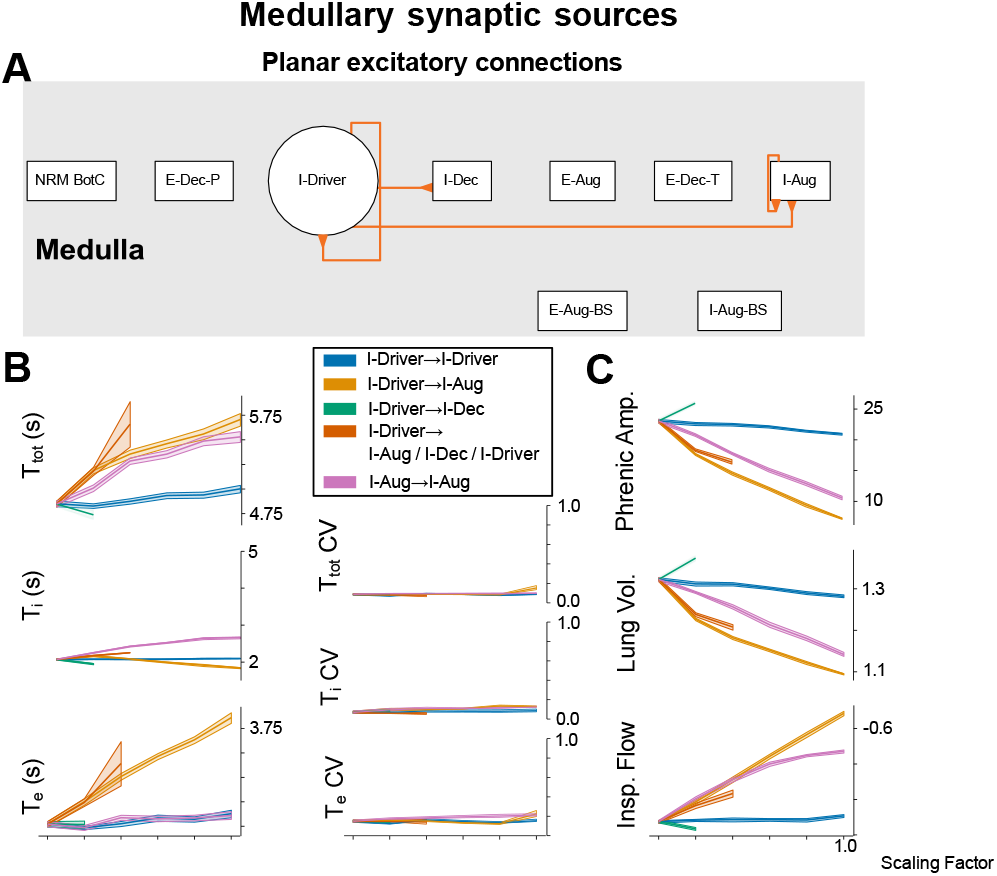
Influence of excitatory synaptic strength on network and biomechanical behaviors through medullary planar connections. **A**, Illustration of the excitatory planar connections from Fig. 1B that are analyzed here. **B**, T_tot_ is respiratory period, T_i_, inspiratory phase duration, and T_e_, the expiratory phase duration. CV is the coefficient of variation for the respective quantities **C**, (top) Maximum phrenic motoneuron amplitude during the inspiratory phase (arbitrary units) (middle) Maximum lung volume during the inspiratory phase. (bottom) Inspiratory flow.

µ-OR agonists have been shown to directly cause Fig. 4A.iii and Fig. 4B.ii in some contexts (Gray et al. 1999; Heinke et al. 2011; Ikoma et al. 2007; Jørgensen et al. 2022; Kim et al. 2024) and Fig. 4B.i may be one mechanism of opioid tolerance (Gillis et al. 2020; Koch and Höllt 2008).

### Alteration of excitability in populations of the upstream core network

We first started by examining the effects of biasing cellular excitability by altering *I*_app_ over the range -10 to +10 pA, where the latter depolarizes neurons. There were 4 conditions, changes in Δ*I*_app_ on the populations of: I-Drivers, I-Drivers + I-Augs, NRM-pons, and all neurons in the simulation for comparison. The results are analogous to the situations demonstrated in Fig. 4Ai-iii.

There were 9 distinct runs of independently generated starting networks (Δ*I*_app_ = 0) for the 4 conditions. Changing Δ*I*_app_ > 0 for all neurons slows the respiratory rhythm (T_tot_) as Δ*I*_app_ >> 0 but also slows the rhythm slightly as inspiratory phase bursts (T_i_) increase when Δ*I*_app_ < 0 (Fig. 5A). If Δ*I*_app_ < 0 falls too low the system loses respiratory activity. As this ceases at Δ*I*_app_ < ∼-2 pA across all networks it shows that the current system is very “stiff” and just on the precipice of cessation if the whole network is seriously perturbed in the hyperpolarizing direction.

For I-Driver + I-Aug perturbations, there is a transient period as Δ*I*_app_ < 0 where the CV of T_tot_, T_i_, and T_e_, increase dramatically compared to similar I-Driver perturbations (Fig. 5A). Curiously, when I-Driver population alone is manipulated the means of both T_i_ and T_e_ roughly track along the same trajectories as I-Driver + I-Aug perturbations. This shows that the I-Aug population is contributing to cycle-to-cycle stability of the respiratory rhythm.

We also modulated NRM-pons neurons to see how they influence overall respiratory activity. While they are non-phasic, the stochasticity of this population’s firing still influences activity in non-intuitive, non-monotonic ways as a function of uniform Δ*I*_app_ (Fig. 5A). At Δ*I*_app_ > 0, breathing became deeper, and lungs are inflated while at Δ*I*_app_ < 0 breaths are shallower (Fig. 5B). Similar trends were found in the more targeted perturbations of I-Drivers and I-Drivers + I-Augs suggesting the NRM-pons neurons are vicariously acting largely through these populations as the connectivity from NRM-pons implies (Fig. 1D).

### Intraplanar medullary synaptic sources affect rhythm generation

The essential elements of the respiratory network are found in the medullary region, so we examined how modulating the maximal excitatory strength 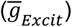 between planar connections in this structure could influence activity (Fig. 6).

Fig. 6A illustrates the subset of anatomically (hypothesized) planar medullary connections from Fig. 1A and 1B and we were focused on the role of excitatory connections. We scaled down the synaptic strength (analogous to Fig. 4B) between the following populations of neurons: I-Drivers → I-Drivers, I-Drivers → I-Augs, I-Drivers → I-Decs, I-Drivers → I-Drivers/I-Augs/I-Decs, and I-Augs → I-Augs (Fig. 6A).

Scaling the strength of recurrent synapses in the I-Driver population (I-Drivers → I-Drivers) resulted in relatively little change in T_tot_, T_i_, or T_e_ (Fig. 6B). Further, perturbation of the strength of these recurrent synapses had little effect on phrenic amplitude, lung volume or peak inspiratory flow (Fig. 6C).

Reducing synaptic strength between the I-Driver and I-Aug populations increased T_tot_ by over 15% and that effect was primarily due to an increase in T_e_ of over 30% (Fig. 6B). There was little effect on T_i_ by this perturbation. Further, there were linear reductions in both phrenic amplitude, lung volume, and peak inspiratory flow (Fig. 6C). Figure 7A shows the spike-time histogram patterns of setting the synaptic strength from I-Driver to I-Aug neurons to 0%. The Phrenic MNs lose robust temporal coherence which explains the reduction in Flow and Lung Volume (compare to Fig. 2).

**Figure 7.**
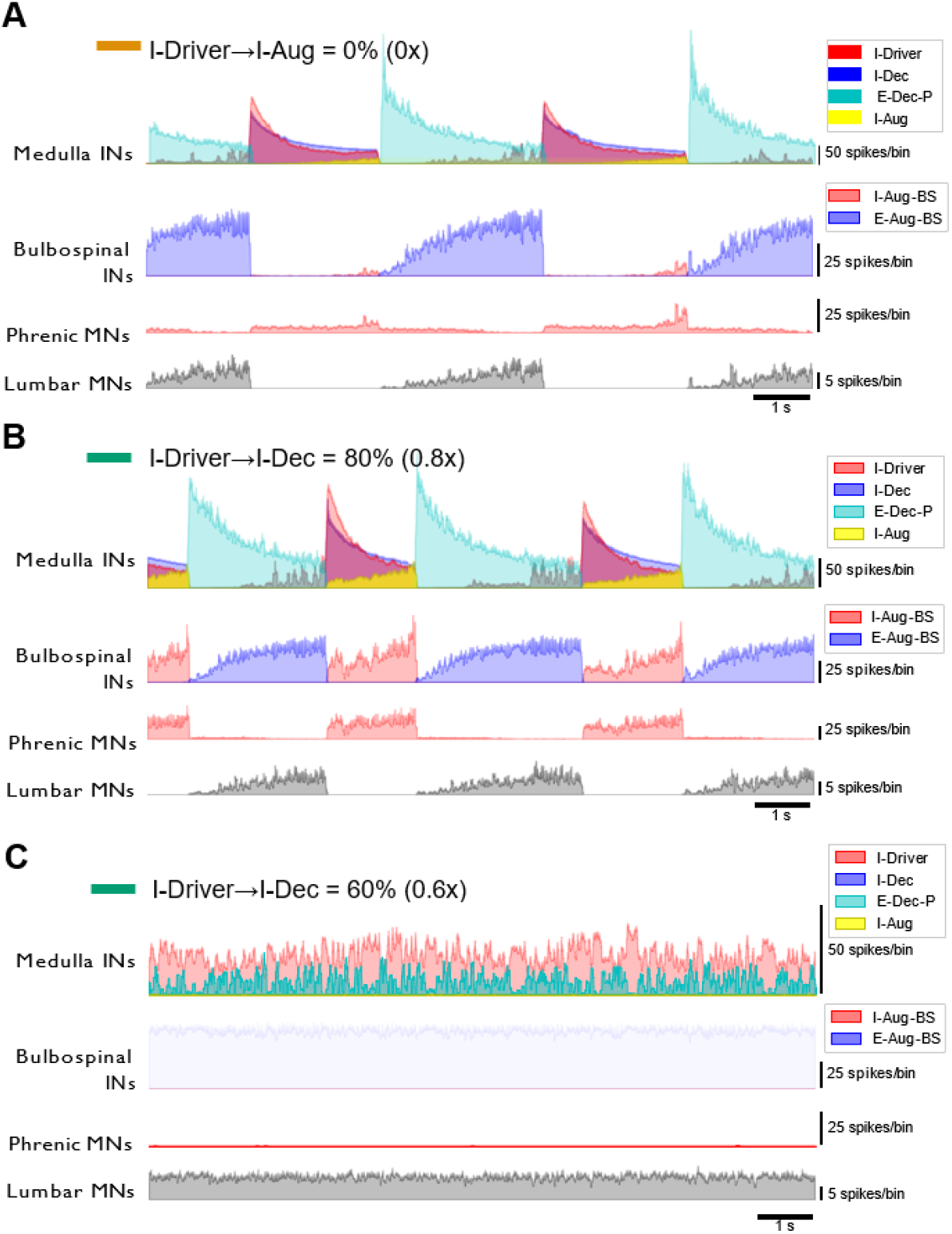
Example activity patterns of key planar excitatory connection perturbations. **A**, Spike-time histogram patterns of the network with I-Driver output to I-Aug neurons at 0% of control. **B**, Spike-time histogram patterns of the network with I-Driver output to I-Dec neurons at 80% of control compared to 60% of control (**C**).

When synaptic strength between the I-Driver and I-Dec populations was reduced, rhythmogenesis and inspiratory motor drive failed after a change between 60-80% (Fig. 6B, C). Figs. 7B and C show examples of firing rate records for medullary and bulbospinal neurons as well as phrenic and abdominal motoneurons during reduction of synaptic strength to 80% (Fig. 7B), and 60% (Fig. 7C) of control for I-Driver to I-Dec synapses.

Simultaneous reductions in the synaptic strength from I-Drivers to other I-Driver neurons, I-Aug neurons and I-Dec neurons resulted in what appeared to be a synthesis of all changes induced by perturbation of excitability for each of the individual populations alone (Fig. 6B, C). As such, simultaneous reductions in synaptic strength by up to 55% increased T_tot_ and T_e_ and decreased phrenic amplitude, lung volume and peak inspiratory flow (Fig. 6B, C). Large reductions in synaptic strength resulted in simulated apnea.

We additionally decreased synaptic strength among recurrent synapses in the I-Aug population alone. Unlike perturbation of synaptic strength among recurrent synapses within the I-Driver population; this action lengthened both T_tot_ and T_i_ by 15-25% with no change in T_e_ (Fig. 6B). Further, phrenic amplitude, lung volume, and peak inspiratory flow were also reduced in a linear manner (Fig. 6C).

### Inhibitory influence of I-Dec hub neurons

Since the I-Dec population of neurons seems to have a dramatic effect on respiratory activity, we decided to also examine the effects of the I-Dec synaptic connections (Fig. 8). This is novel in comparison to the previous figures in that we were probing the influence of inhibitory synapses.

**Figure 8.**
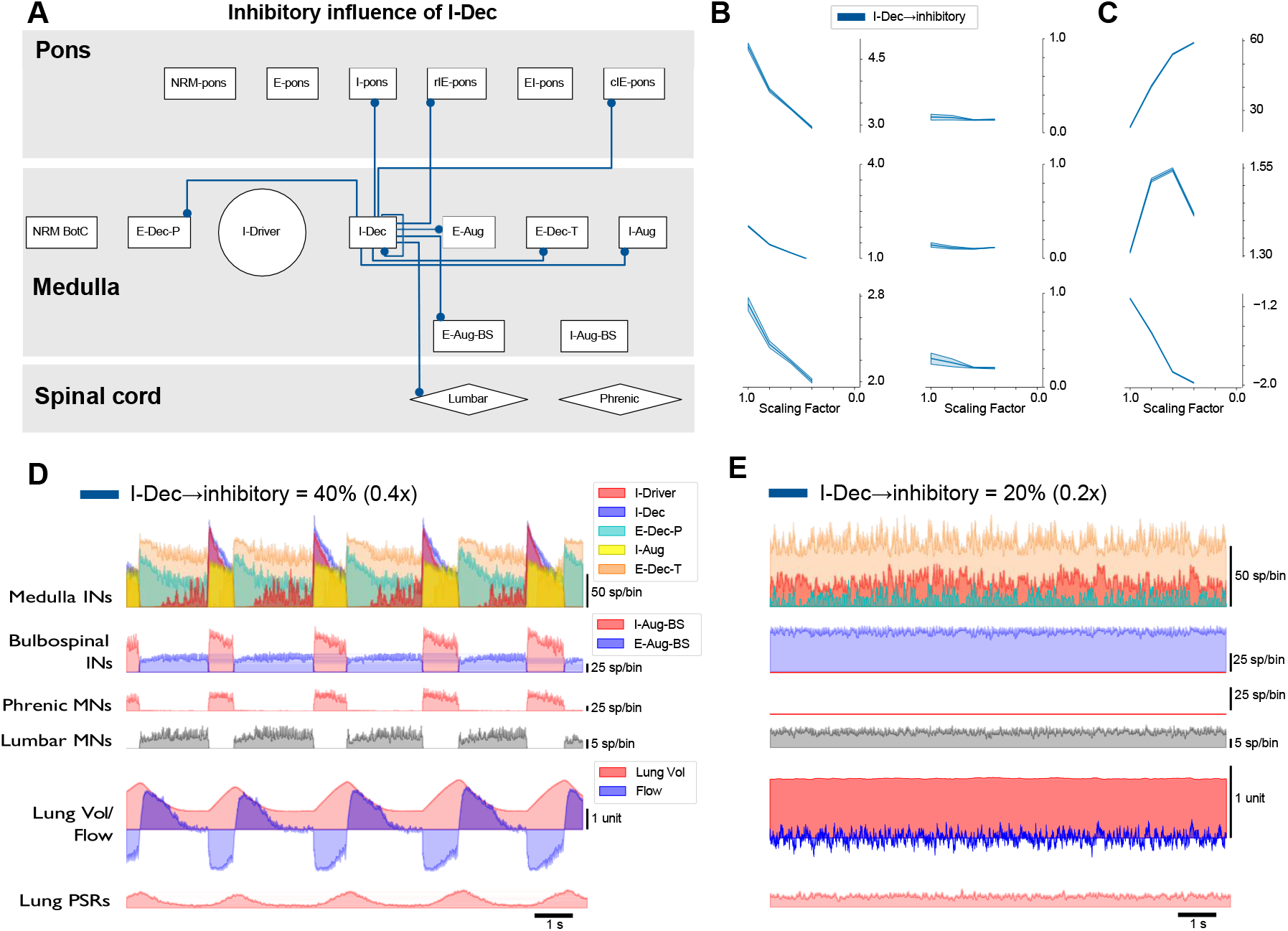
Influence of inhibitory synaptic strength on network and biomechanical behaviors through I-Dec hub connections. **A**, Illustration of the inhibitory connections from I-Dec neurons in Fig. 1A that are collectively altered here. **B**, T_tot_ is respiratory period, T_i_, inspiratory phase duration, and T_e_, the expiratory phase duration. CV is the coefficient of variation for the respective quantities. **C**, (top) Maximum phrenic motoneuron amplitude during the inspiratory phase (arbitrary units). (middle) Maximum lung volume during the inspiratory phase. (bottom) Inspiratory flow. **D**, The spike-time histogram patterns of the network with I-Dec inhibitory output at 60% of control compared to 40% of control (**E**).

Perturbing all the inhibitory connections from the I-Dec population (Fig. 8A) causes the respiratory behavior to drop (Fig. 8B/C) because these neurons are the hub of our system with connections to 19 of the other 46 populations of neurons (10 inhibitory connections shown). In general, as the maximal inhibitory strength 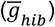 decrease the T_i_, T_e_, and T_tot_ get shorter and these quantities get more regular (Fig. 8B), and the phrenic activity monotonically increases (Fig. 8C). When 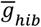 is 60% of control, Fig.8D demonstrates hyperpnea-like activity with intense inspiratory/expiratory activity and large lung inflations/deflations before the rhythm goes out in Fig. 8E when 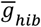 drops to 55% and lower.

### Descending pontine synaptic sources affect burst patterning

Finally, we also looked at how perturbing 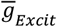 between NRM and I-Aug or I-Driver neurons affected the overall breathing pattern. Decreasing the excitatory connections from the pontine synaptic sources (Fig. 9A) led to disordered and inconsistent T_i_, T_e_, and T_tot._ production (Fig 9B). When 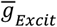 is blocked, phrenic activity does increase for the I-Aug + I-Driver and I-Driver connections. However, it decreases for the I-Aug connections and remains unchanged for the I-pons connections (Fig. 9C). Blocking this descending pontine transmission led to cluster-like breathing (Fig. 10).

**Figure 9.**
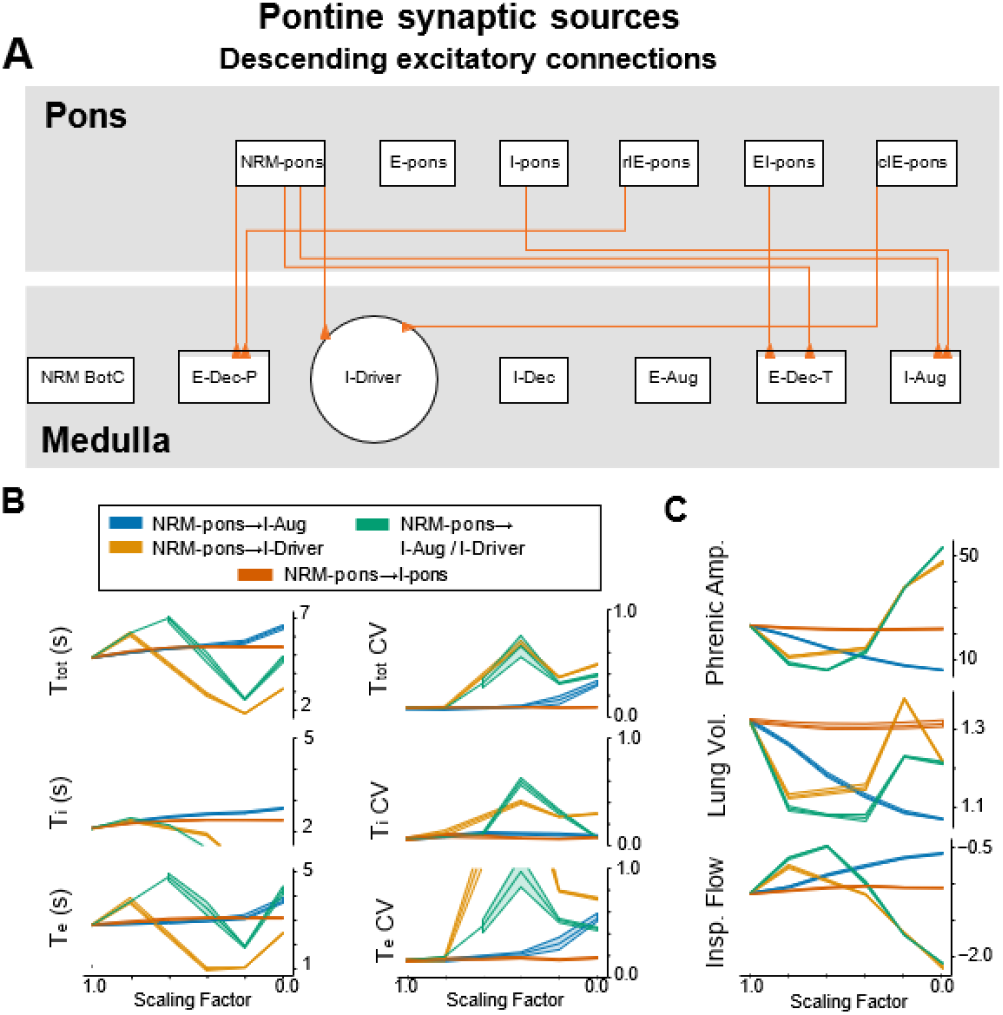
Influence of excitatory synaptic strength on network and biomechanical behaviors through descending pontine-medullary connections. **A**, Illustration of the descending excitatory connections from pontine non-respiratory modulated neurons (NRM-pons), phase spanning neurons (rIE-pons, cIE-pons, EI-pons), and inspiratory neurons (I-pons). **B**, T_tot_ is respiratory period, T_i_, inspiratory phase duration, and T_e_, the expiratory phase duration. CV is the coefficient of variation for the respective quantities. **C**, (top) Maximum phrenic motoneuron amplitude during the inspiratory phase (arbitrary units). (middle) Maximum lung volume during the inspiratory phase. (bottom) Inspiratory flow.

**Figure 10.**
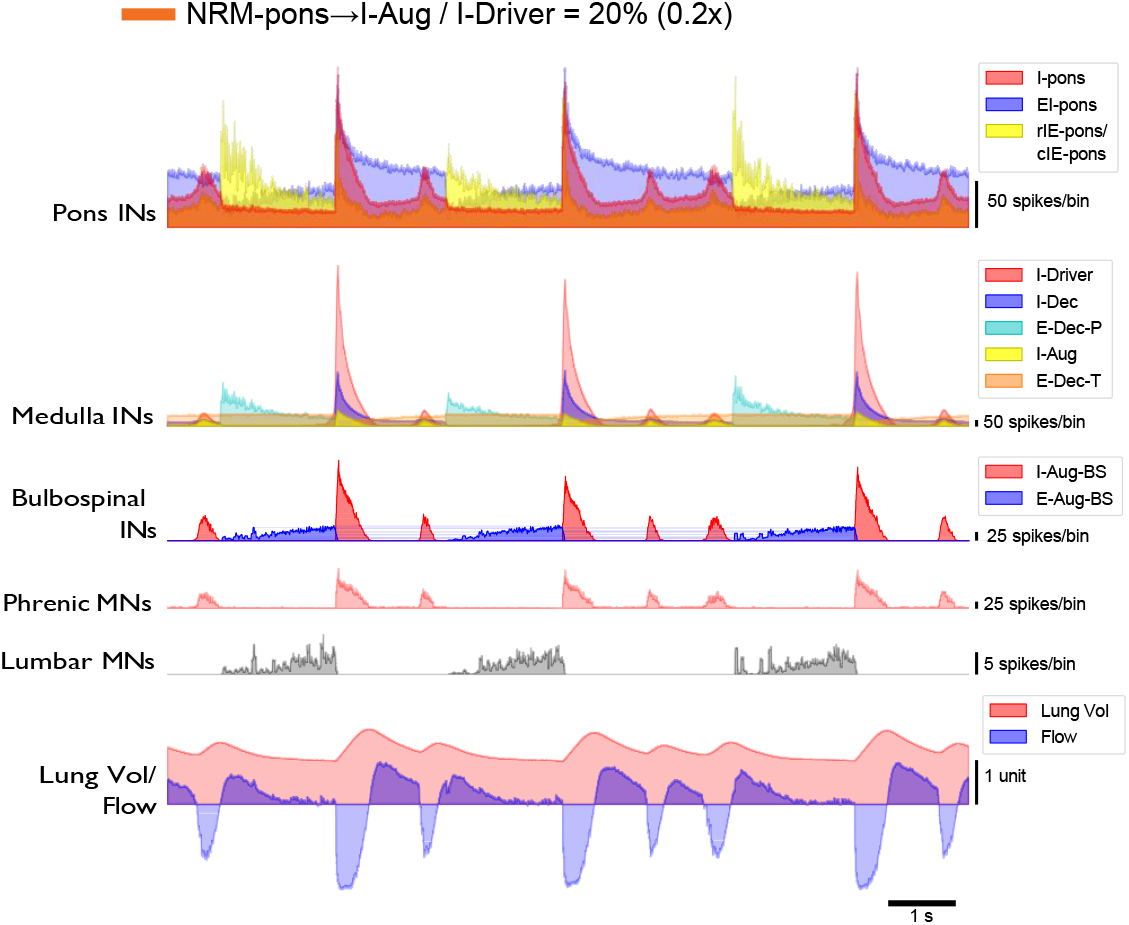
Example activity patterns of descending excitatory connection perturbations from NRM-pons neurons. The network pattern with NRM-pons excitatory output at 20% of control projecting to I-Aug and I-Driver populations resulting in clustered-like bursts in inspiratory activity.

## DISCUSSION

The effects of opioids on respiratory function in experimental conditions remains a critical research priority. There are many effects that opioids pose on respiratory function in experimental and clinical conditions (e.g., decreases or abnormal function in chest wall compliance, tidal volume, respiratory rate, etc.). This has led to much investigation that has focused on the impact of different opioid agonists on respiratory depression (Skulsky et al. 2007). The advantages computational models present are the ability to manipulate parameters that are experimentally inaccessible and make predictions about the resulting effects. The current study investigated the simulated responses of activating µ-ORs within a computational model of the pontomedullary respiratory network to better understand the neural mechanisms contributing to OIRD. Since morphine, codeine, and similar drugs, have multiple side effects beyond just activating µ-ORs within the brainstem (Dahan 2007; Simera et al. 2010; Tomazini Martins et al. 2018), our model is best interpreted to most closely reproduce the highly specific µ-OR ligand fentanyl and its effects on respiratory activity, rhythm changes, current alterations, and spike burst changes within the brainstem network (Lalley 2003; Shen et al. 2022). It is important to note, that Figures 1 and 2 depict eupneic conditions within the model.

### Simulated opioid effects on respiratory activity

Previous computational models have perturbed the connection of fictive medullary neurons within the brainstem (Chou et al. 2024; Shevtsova et al. 2011; Magosso et al. 2004). The advantage of the current study, with the employed joint neuronal network-biomechanical model, is that we examined these factors at a biomechanical level and in the context of a broader brainstem neuronal network. The strength of the network’s connectivity is an important parameter that affects the neural breathing patterns, and the model is generally inhibited when perturbed by opioids, specifically simulated fentanyl. While decreasing the connection strength or explicitly hyperpolarizing member populations, our results indicate that breathing patterns were significantly affected and had an overall inhibitory effect (see Figs 8 and 10). OIRD is characterized by a decrease in respiratory rate and irregular breathing frequencies and, at high doses, apnea. This has been attributed to opioids activating G-coupled proteins through µ-ORs which hyperpolarize cells through G-protein-gated inward rectifying K^+^ (GIRK) channels (Montandon et al. 2016a, 2016b). Furthermore, the current model incorporates neuron populations (i.e., NRM (non-respiratory modulated), tonic-expiratory motoneurons, and interneurons) at a broader brainstem level than previous models. Previous neural recordings have reported the importance of interneurons and their prevalence through the respiratory network (Lindsey et al. 2012; Segers et al. 2008, 2015). Therefore, including these populations (i.e., the interneurons between the I-Driver and bulbospinal population) within the current simulations is essential in understanding the brainstem network’s respiratory dynamics.

Researchers have reported decreased respiratory rates when opioids were directly applied to the ventrolateral medulla or systemically injected (Montandon et al. 2016a, 2016b; Suzue 1984). For example, when DAMGO (d-Ala2, *N*-MePhe4, Gly-ol]-enkephalin), was applied to the ventrolateral medulla and presumably activating local µ-ORs, it reduced the respiratory rate of mice, but did not affect the diaphragm amplitude in GIRK2^−/−^ mice (Montandon 2022). In the same study, a moderate intramuscular injection of fentanyl was provided to the GIRK2^−/−^ mice, and only a slight depression in diaphragm amplitude was observed. In a complementary study, systemic administration of fentanyl reportedly decreased respiratory rate, yet had no effect on diaphragm amplitude (Montandon et al. 2016a, 2016b).

### Simulated opioid effects on respiratory motoneuronal bursts

Within our simulations, altering the synaptic strength of the pontomedullary inspiratory neurons affected spike burst durations (T_i_) until the respiratory activity was extinguished (Figs. 7A, 7B, and 7C). More specifically, as synaptic conductance was systematically decreased, the I-Driver to I-Aug connection resulted in a prolonged inspiratory burst with a ramping effect. As the I-Driver to I-Dec connection within the model was perturbed, simulations demonstrated a profound reduction in phrenic motor bursting; indicating that the I-Dec population may serve as an “off-switch” population within the network that are opioid sensitive (Fig. 7). These simulation results are supported by previous findings. Specifically, the administration of opioids has been shown to affect the burst duration of respiratory motor units up to the point of respiratory arrest (Lalley 2006). As discussed above, one mechanism opioids likely perturb respiratory patterns is through cell hyperpolarization, which in turn, affects the spiking activity of respiratory neurons (Fig. 4A). In our simulations, modulating an injected bias current (*I*_app_) within the core medullary populations (Fig. 5) affected spike burst durations (T_i_) until the respiratory activity was extinguished at larger hyperpolarizing *I*_app_. Therefore, within our modeling efforts, modifying the injected current qualitatively reproduces *in vivo* effects of OIRD.

Lalley (2003) investigated the intravenous effects of fentanyl in vagotomized adult cats while recording individual neurons. He reported prolonged discharges that induced tonic firing of bulbospinal expiratory neurons (like our model’s E-Aug-BS population) that were correlated with a reduced hyperpolarization of synaptic drive potentials. Lalley suggested that this result may have been explained by the decrease in the duration of the inspiratory phase observed at certain dose-responses of fentanyl (Lalley 2003). He further interpreted lower doses of fentanyl to have a similar effect on vagal post-inspiratory motoneurons which led to “sparse, low-frequency” discharges which suggests that fentanyl regulates bulbospinal, and motoneurons, presynaptically at different dose-dependent responses (Lalley 2003; Lalley and Mifflin 2017). Our model simulations also add plausibility to their conclusions, especially those represented by a combination of Fig. 4Aiii and Fig. 4Bii. These simulation results further support the findings of Lalley and Mifflin (Lalley and Mifflin 2017). Their findings postulated that µ-opioid agonists directly affect the controlling and timing of burst and oscillation patterns of bulbospinal and vagal motoneurons, which also have a direct effect on respiratory muscle force. The plausibility of their conclusions is supported by our simulated alterations in synaptic conductance between the I-Driver and I-Aug and the I-Driver and I-Dec motoneuron populations.

### Effects of changes in membrane current

An important consideration for this model is that the I-Driver population are fundamentally essential for any kind of respiratory patterning under our simulated conditions. All members of that population burst based on a slowly-inactivating persistent sodium current (*I*_*NaP*_) (Breen et al. 2003; Butera et al. 1999) which has been recently shown to be inessential for I-Driver-like activity (da Silva et al. 2023). For these simulations, there is no salient difference in what bursting mechanism we choose for the I-Driver population as we are interested in network effects as emphasized by the depiction in Fig. 3. However, when 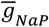 was altered, this led to the cessation of the network rhythm (Fig. 3D). Essentially by decreasing *I*_*NaP*_ the cells broadly hyperpolarize, and the network aborted the respiratory rhythm. However, as we showed in Figure 3, modulating the parameters determining the qualities of 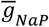 on individual I-Driver neurons has a dichotomous effect versus how the more expansive multi-population network behaves.

Previous findings by Baertsch and coworkers (Baertsch et al. 2021). Following the administration of intravenous fentanyl, researchers documented changes in current and discharge properties of neurons, which in turn, caused changes in overall timing and frequency of respiratory behavior (Baertsch et al. 2021; Lalley and Mifflin 2017).

### Limitations of the model

While we incorporate vagal afferent feedback as well as respiratory mechanics into our model, we have not simulated central chemoreceptors. Therefore, our simulations emulate *in vivo* models in which CO_2_ is clamped by mechanical ventilation. In the non-mechanically ventilated condition, CO_2_ would rise with the depressant actions of opioids and compensate by increasing respiratory drive through the hypercapneic ventilatory response (HCVR), thereby blunting the actions of these drugs on the respiratory control system (Baldo 2021; Pattinson 2008; Paul et al. 2021). However, CO_2_ does not rise by large amounts because the HCVR is open loop.

### Physiological and clinical implications

A crucial motivation for this kind of modeling study is that experimental studies of this kind are currently unfeasible. Here, we are delineating neuronal populations of interest both anatomically and, more importantly, functionally, because the respiratory network is distributed across much of the brainstem (Segers et al. 2008). Thus, providing a systematic assessment of how the opioid agonist, fentanyl, affects the inspiratory motoneurons within the preBötzinger and pontine circuitry. Secondly, the model allowed our team to evaluate the antecedent pathways within the pontomedullary network that may be µ-opioid sensitive. Lastly, we were able to identify a neuron population within the computational network that is opioid sensitive and is supported by previous *in vivo* findings (Lalley and Mifflin 2017). With advancing genetic technologies these avenues may be more closely explored but challenges remain and may require higher-order intersectional approaches than what is currently common.

In the case of anatomical specificity, approaches such as viral injections into specific locations can partially address these issues (Liu et al. 2022; Varga et al. 2020). This may be especially the case when expression from these injections is conditioned on specific gene promoters (Nectow and Nestler 2020). However, specifying genetic tools to firing pattern is nebulous for the most part. A combination of ion channel expression, endogenous Ca^2+^ buffers such as parvalbumin (Alheid et al. 2002), synaptic partners, or constitutively expressed transcription factors (Bachmutsky et al. 2020; Sun et al. 2019), may provide a means of intersectionally subdividing certain populations given a certain neuronal population’s “fingerprint” of multiple distinguishing criteria but that remains beyond the scope of the present study.

### Summary

Opioids are clinically used for their analgesic effects perioperatively; however, their use can lead to respiratory depression and the disfacilitation of airway protective mechanisms. The early detection of respiratory suppression allows clinicians to make life-saving decisions and avert the catastrophic consequences of OIRD. The overall results of our modeling efforts indicate that the joint neuronal-biomechanical model employed in the current study demonstrated overall inhibition, frequency alterations, spike burst changes, and timing changes, which are supported by the different perturbations observed within *in vivo* data that employs the µ-OR agonist fentanyl. The proposed model is an excellent tool that lends itself to answering questions that persist within the opioid crisis, specifically revolving around OIRD.

## Supporting information

Supplemental Table of Target Populations

## DATA AVAILABILITY

The datasets generated and analyzed during the present study are available from the corresponding author on reasonable request.

