## Supplemental Table of Target Populations for "Modeling Insights into Potential Mechanisms of Opioid-Induced Respiratory Depression within Medullary and Pontine Networks"

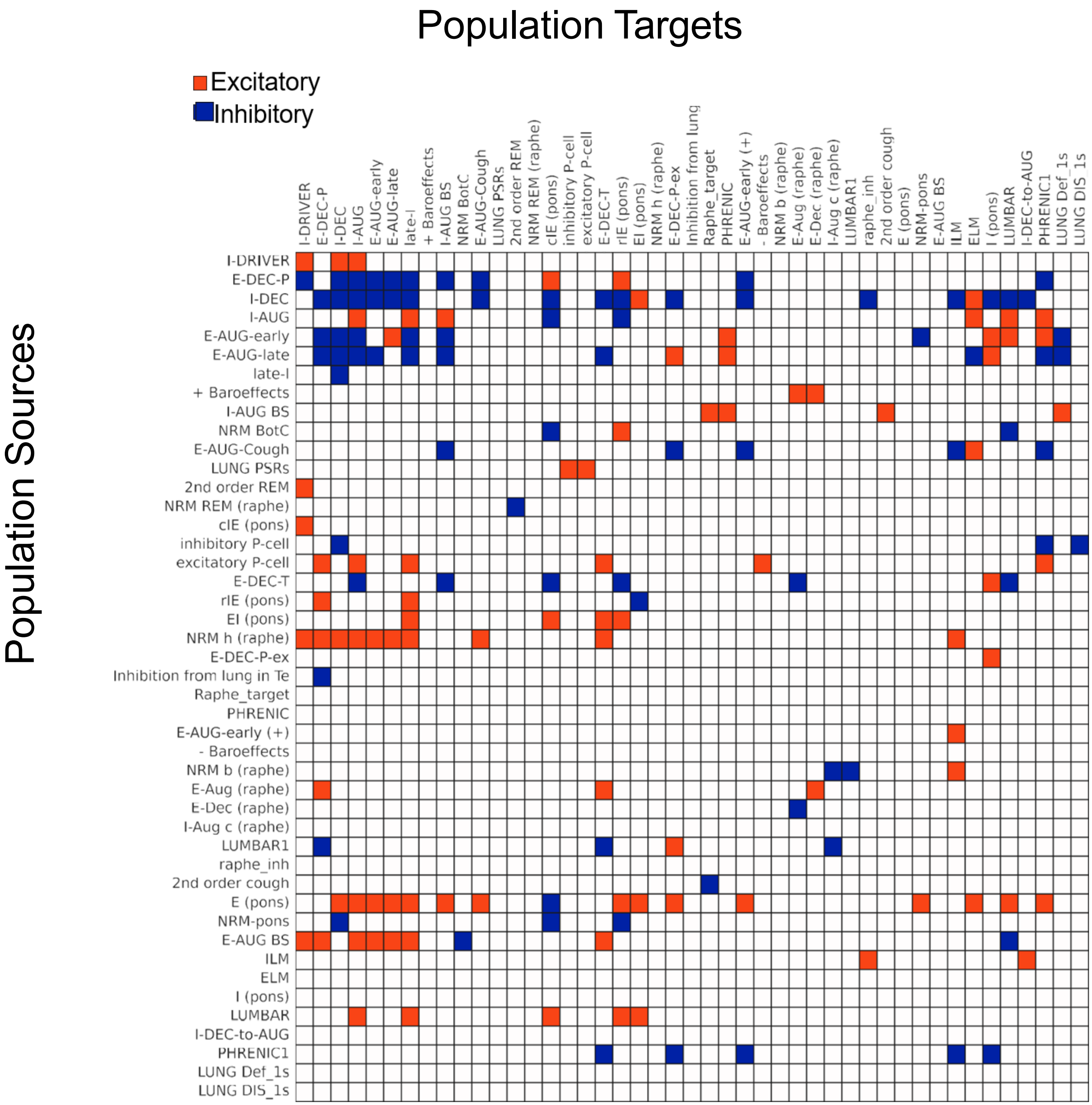


**Figure S1. Global connectivity adjacency matrix.** Identifies which source cell populations (rows) connect to which target cell populations (columns) with excitatory (orange), inhibitory (blue), or the absence of connections (white).
